# Naturally occurring CodY variants alter ligand binding, DNA target affinity, and virulence in *Clostridioides difficile*

**DOI:** 10.1101/2025.11.06.687015

**Authors:** Md Kamrul Hasan, Sourav Roy, Revathi Govind

## Abstract

*Clostridioides difficile* is an important nosocomial pathogen that has been classified as an “urgent threat” by the CDC. Antibiotic use is the primary risk factor for the development of *C. difficile*-associated disease because it disrupts healthy protective gut flora and enables *C. difficile* to colonize the colon. *C. difficile* damages host tissue by secreting toxins and disseminates by forming spores. Nutrient availability and other environmental factors greatly influence toxin production in *C. difficile*. CodY is a global transcriptional regulator that coordinates metabolism and virulence in Gram-positive pathogens in response to nutrient availability, primarily through sensing branched-chain amino acids (isoleucine, leucine, valine; ILV) and GTP. In *C. difficile*, CodY indirectly represses toxin production by inhibiting transcription of the positive regulator *tcdR* and influences sporulation through unknown mechanisms. Here, we characterized two naturally occurring CodY variants, CodY(Y146N) and CodY(V58A). The Y146N substitution is located near the GTP-binding pocket, while V58A is near the ILV-binding site. Ligand binding assays revealed GTP binding ability of CodY(Y146N) is severely compromised. Electrophoretic mobility shift assays (EMSAs) demonstrated that ligand binding differentially influenced promoter binding; in the presence of GTP, CodY(Y146N) bound to the *tcdR* promoter less efficiently than CodY(WT). In *C. difficile*, production of either variant resulted in reduced repression of toxin production compared to CodY(WT). Subsequent *in vivo* experiments in a hamster infection model showed that strains producing CodY(Y146N) or CodY(V58A) were significantly more virulent than the CodY(WT) producing strain. These findings demonstrate that a single amino acid change in this global regulator can alter its ligand affinity and promoter-binding properties to potentially rewire the gene regulatory networks to enhance the pathogenic potential in *C. difficile*.

**Importance:** *C. difficile* has been recognized as an important nosocomial pathogen that causes diarrheal disease as a consequence of antibiotic exposure. CodY controls the expression of numerous metabolic and virulence genes. In this study, we have characterized two CodY variants and have demonstrated that even a single residue change in important domains can affect its function and have implications on bacterial virulence.

## Introduction

*Clostridioides difficile* is a Gram-positive, spore-forming anaerobe and a major cause of antibiotic-associated diarrhea and colitis in hospitalized patients. In the United States alone, *C. difficile* infections (CDI) are associated with ∼20,000 deaths annually and incur about 4.8 billion USD in Healthcare costs (Dubberke and Olsen, 2012; Lessa *et al*., 2015). Disease severity results primarily from the production of two large clostridial toxins, TcdA and TcdB, which disrupt host epithelial barriers, trigger inflammation, and cause tissue damage (Voth and Ballard, 2005; Smits *et al*., 2016). The regulation of toxin synthesis and sporulation is tightly linked to nutrient availability in the host environment (Dineen *et al*., 2010).

CodY functions as a global transcriptional regulator responsible for nutrient sensing and is highly conserved among low G+C Gram-positive bacterial pathogens such as *Bacillus anthracis, Staphylococcus aureus, Streptococcus mutans,* and *Listeria monocytogenes* (Sonenshein, 2005), (Stenz *et al*., 2011). In nutrient-rich conditions, CodY represses the expression of the genes it regulates. This repression occurs through CodY binding near the promoter regions of target genes at a specific 15-nucleotide consensus sequence, AATTTTCWGAAAATT, where W represents either adenine (A) or thymine (T) (Belitsky and Sonenshein, 2008). The interaction of CodY with DNA is further strengthened by its binding with branched-chain amino acids, specifically isoleucine, leucine, and valine (ILV), as well as GTP, either independently or in combination (Shivers and Sonenshein, 2004; Handke *et al*., 2008; Brinsmade and Sonenshein, 2011). The presence of ILV and GTP modulates CodY’s DNA-binding capability, enabling the protein to adapt to the cell’s nutritional status. Structural studies have shown that CodY senses ILV and GTP through distinct ligand-binding sites, leading to conformational changes that modulate its DNA-binding activity (Levdikov *et al*., 2006; Levdikov *et al*., 2017). Under nutrient-deficient conditions, the insufficient availability of ILV and GTP alleviates CodY-mediated repression of the target genes (Ratnayake-Lecamwasam *et al*., 2001; Brinsmade and Sonenshein, 2011). CodY is recognized for regulating genes associated with carbon overflow metabolism, the Krebs cycle, synthesis of certain amino acids, genetic competence, and the uptake and catabolism of amino acids, peptides, and sugars(Sonenshein, 2007a).

In *C. difficile*, the genes *tcdA* and *tcdB*, which encode toxins A and B, are located within the pathogenicity locus (PaLoc) that also contains *tcdR*, *tcdE*, and *tcdC* (Cohen *et al*., 2000). The synthesis of toxins is augmented as *C. difficile* cells transition to the stationary phase and is significantly modulated by nutrient availability. A definitive association between nutrient limitation and the expression of toxin-related genes has been established through the activity of CodY, which represses the transcription of all genes within the PaLoc, including *tcdR*, the toxin-specific sigma factor that directs the transcription of *tcdA* and *tcdB* (Dineen *et al*., 2007; Dineen *et al*., 2010). CodY also represses sporulation initiation, possibly through regulation of *spo0A* activity, although the underlying mechanisms remain unclear (Nawrocki *et al*., 2016).

Analysis of *C. difficile* genome sequences has revealed multiple naturally occurring *codY* alleles. Thousands of *C. difficile* strains have been sequenced so far by various research groups around the world, and their genomes are readily available to the research community for analysis through the NCBI database. Among these, most of the *C. difficile* strains carry the *codY* gene, which we depict as *codY(*WT) in this work. BLASTP analysis using the CodY(WT) sequence revealed a few variants among *C. difficile* CodY. These include alleles with the following mutations: L55M, A46S, G152S, E89Q, T18K, and V58A, and Y146N. Among these, two variants, Y146N and V58A, are of particular interest because the substitutions are located near the predicted GTP- and ILV-binding pockets, respectively. The CodY(Y146N) allele has been identified in one of the derivatives of the epidemic ribotype 027 strain R20291 (Monteford *et al*., 2021), while CodY(V58A) occurs in more than 170 clinical isolates spanning diverse ribotypes, including ribotype 078, which is associated with animal reservoirs and emerging human infections. We hypothesized that these naturally occurring substitutions alter CodY’s ligand-binding properties, DNA target specificity, and regulatory activity, with potential consequences for *C. difficile* physiology and virulence.

Here, we characterized CodY(Y146N) and CodY(V58A) using biochemical, genetic, and *in vivo* approaches. Ligand binding assays demonstrated that the Y146N substitution affected GTP binding, whereas V58A reduced ILV binding. Electrophoretic mobility shift assays (EMSAs) revealed altered promoter-binding responses to ligands. Allele replacement in an isogenic *C. difficile* background showed that both variants reduced repression of toxin production. In a hamster infection model, CodY(Y146N) and CodY(V58A) producing strains exhibited enhanced colonization and virulence compared to the CodY(WT) strain. These findings indicate that single amino acid changes in CodY can rewire regulatory networks and enhance pathogenic traits in *C. difficile*.

## Materials and methods

### Bacterial strains and growth conditions

Bacterial strains and plasmids used in this study are listed in Table S1. *C. difficile* strains were grown anaerobically (10% H2, 10% C02, and 80% N2) in Ty (tryptose yeast extract) agar or broth as described previously (Girinathan *et al*., 2017; Girinathan *et al*., 2018). Kanamycin (Kan; 50 µg/ml), Cefoxitin (Cef; 25 µg/ml), and Thiamphenicol (Thio; 15 µg/ml) were added to culture medium whenever necessary. Guanosine (100 µM) and/ or ILV (isoleucine, leucine, and Valine; 10 mM) were added to TY broth to measure the effect of CodY-ligands on toxin production. Escherichia coli S17-1 (6) strains carrying specific Clostridial shuttle plasmids were cultured aerobically in Luria-Bertani (LB) broth and supplemented with Chloramphenicol (25µg ml-1).

### Cloning, expression, and purification of recombinant CodY proteins in *E. coli*

The full-length *codY* genes encoding CodY(WT), CodY(V58A), and CodY(Y146N) were cloned and overexpressed in Rossetta *E. coli* cells using pET16B expression system. The *CodY(WT)* and *CodY(Y146N)* genes were PCR amplified using chromosomal DNA templates of *C. difficile* strains UK1 and R20291, respectively. The PCR fragments were then digested with Xho1 and BamHI and were cloned into pET16B digested with the same. The resulting plasmids pRG359 and pRG361 (S1 Table) were then transformed into *E. coli* Rossette competent cells for the protein expression. Site-directed mutagenesis was used to introduce the V58A mutation in the pRG361 to generate pRG367. The *E. coli* strains carrying the *codY* constructs were grown at 37°C in LB medium containing chloramphenicol (25µg ml-1) and ampicillin (100µg ml-1). Protein expression was achieved by inducing with 1mM IPTG at 17°C overnight. Cells were harvested by centrifugation, and the His-tagged proteins were purified by nickel-affinity chromatography (Sigma) in buffer containing 50mM sodium phosphate (pH 8.0), 0.3M NaCl, and 10mM imidazole. The Nickel-bound proteins were eluted with a linear gradient of imidazole (50-500 mM). The purified proteins collected were analyzed by SDS-PAGE for purity and were then concentrated by passing through Amicon membranes with a cut-off value of 5KDa and equilibrated in phosphate buffer with 20% glycerol before storage at -80 °C.

### Size Exclusion Chromatography

The oligomeric state of CodY and its mutant variants was analyzed by size-exclusion chromatography (SEC). Purified His-tagged CodY(WT), CodY(V58A), and CodY(Y146N) proteins were dialyzed against SEC buffer containing 20 mM Tris-HCl (pH 8.0), 150 mM NaCl, and 5 mM MgCl₂. Each sample (500 µL, 2–5 mg/mL) was loaded onto a Superdex™ 200 Increase 10/300 GL column (Cytiva) equilibrated with the same buffer and operated at a flow rate of 0.5 mL/min using an ÄKTA Pure chromatography system. Protein elution was monitored at 280 nm, and the elution volumes were recorded to estimate the apparent molecular weights of the proteins. The column was calibrated with a set of molecular weight standards (thyroglobulin, γ-globulin, ovalbumin, myoglobin, and vitamin B_12_ Bio-Rad) under identical conditions. The apparent molecular weights of CodY and its variants were calculated based on a standard curve generated by plotting the logarithm of the molecular weights of the standards against their elution volumes.

### Ligand Binding Assays

Purified CodY proteins (50 µM) were incubated with a 5-fold increasing gradient of biotin-GTP (NEB labs) concentrations (5–125 µM) in binding buffer (20 mM Tris-HCl, pH 8.0; 1 mM EDTA; 0.5 mM dithiothreitol [DTT]; 250 mM KCl; 2 µg/mL bovine serum albumin) for 20 minutes. Ten microliters of the reaction mixture were spotted onto pre-activated PVDF membranes and allowed to air dry. The membranes were washed three times with PBS-T and incubated overnight at 4 °C with streptavidin-HRP conjugate antibody (1:5000 dilution). The dot blot was developed using a chemiluminescent HRP substrate and imaged with an Azure 300 Imaging System. For the competition assay, 50 µM CodY (WT) was incubated with 25 µM biotin-GTP along with increasing concentrations of unbiotinylated GTP (5–125 µM). The dot blot procedure was performed as described above. For the radioactive CodY GTP-binding assay, purified CodY proteins at varying concentrations (5 µM–40 nM) were incubated with [γ-³²P]GTP (PerkinElmer Life Sciences) in GTP-binding buffer, followed by UV-mediated cross-linking. Reaction samples were resolved on 12% SDS-PAGE and visualized by autoradiography using a Typhoon imager.

For the leucine-binding assay, purified CodY proteins (20 µg) were incubated with [³H]-leucine, followed by UV-mediated cross-linking. Proteins were TCA-precipitated, and radioactivity in the precipitated proteins was measured using a Beckman Coulter LS6500 scintillation counter.

### Generating *C. difficile* strains expressing *codY* variants

The *codY* alleles were introduced into *C. difficile* genome using the pMSR0 plasmid-based allele exchange method (Peltier *et al*., 2020). Briefly, we cloned *codY*(Y146N), *codY*(V58A), and *codY*(WT) genes flanked with 500 bps of upstream and downstream DNA in the pMSR0 plasmid vector (S1 Table). These plasmid constructs were then introduced into *C. difficile* UK1::*codY* mutant where the original *codY* gene was interrupted by a group II intron(Daou *et al*., 2019a) .*C. difficile* transconjugants were selected on TY agar supplemented with cefoxitin, D-cycloserine, and thiamphenicol. Single cross-over integrants were selected based on the size of the colonies after re-streaking of the transconjugants on BHI with thiamphenicol. Colonies that underwent the second recombination event were then selected on BHI plates containing anhydrotetracycline (ATc: 100 ng/ml). Thioamphenicol-sensitive colonies were then tested by Western blot using CodY-specific antibodies.

### Toxin ELISA

Relative quantification of cytosolic toxins in the *C. difficile* cultures grown in the TY medium was performed as described previously with slight modifications (Girinathan *et al*., 2017). In brief, bacterial cells harvested from 12 h old *C. difficile* cultures were resuspended in 200 µl of sterile PBS and were sonicated to extract cytosolic proteins. One 100 µg of cytosolic proteins from each sample was then used to measure the relative toxin levels using the *C. difficile* Premier Toxin A B ELISA kit from Meridian Diagnostics Inc (Cincinnati, OH), following the manufacturer’s instructions.

### Intracellular c-di-GMP measurement

*C. difficile* cells grown overnight in the TY agar medium were scooped from agar plates and resuspended in 1mL PBS buffer at an OD600 of 2.0. Pellets of cells were washed twice with PBS and resuspended in 200µL PBS. The cells were lysed by sonicating on ice and centrifuged. The cytosolic contents collected were used to measure intracellular c-di-GMP level using Lucerna Cyclic-di-GMP Assay Kit following the manufacturer’s recommended protocol. Briefly, 20µL of the cytosol was incubated with the regents supplied (c-di-GMP sensor and the fluorophore), and the resulting fluorescence signal was measured using GFP/FITC filter sets on Biotek Synergy Plate Reader. Intracellular c-di-GMP concentration was calculated using a standard curve created with known concentrations of pure c-di-GMP.

### Western Blot Analysis

C. *difficile* cells for western blot analysis were harvested and washed in 1x PBS solution before suspending in sample buffer (Tris 80mM; SDS 2%; and Glycerol 10%) for sonication. Whole cell extracts were then heated at 100^°^C for 7 min and centrifuged at 17,000 g for 1 min, and the proteins were separated by SDS-PAGE and electroblotted onto PVDF membrane. Immobilized proteins in the membranes were then probed with CodY-specific antibodies at a dilution of 1:10,000. The blot was subsequently probed with HRP-conjugated secondary antibodies at a dilution of 1:10,000. Immuno-detection of proteins was performed with the Azure 300 imaging system.

### Electrophoretic Mobility Shift Assay (EMSA)

For gel mobility shift assays, *tcdR* upstream DNA with the predicted CodY binding sequence was amplified using primers ORG719 and ORG720, and the PCR product of ∼350 bps was cloned into the pGEMT cloning vector. The DNA region was excised using EcoRI and was then radiolabeled with [α-^32^P] dATP-6000 Ci/mmol (PerkinElmer Life Sciences) using the Klenow fragment of DNA polymerase. The DNA-protein binding reaction was carried out by mixing labeled DNA with increasing amounts of purified CodY proteins in 20µl binding buffer [10mM Tris pH 8.0, 50mM KCl, 50µg BSA, 0.05% NP40, 10% Glycerol, and 250ng of calf thymus DNA]. The ILV (10mM each of isoleucine, leucine, and valine), or 2mM GTP, or both, were added to the binding reactions when needed. The binding reactions were performed at room temperature for 30 min and were stopped by adding 5µl of gel loading buffer. Reactions were loaded onto a 6% native polyacrylamide gel in 1XTBE (Tris/Borate/EDTA) and subjected to electrophoresis at 100 V for 45 minutes. If 10mM ILV was present in the binding reaction, the same concentration of ILV was also added to the electrophoresis buffer. Gels were then dried under vacuum for 1hr and the autoradiography was performed using Typhoon with Molecular Dynamics Phosphor-Imager technology.

### Hamster model to evaluate *C. difficile* virulence

Male and female Syrian golden hamsters were used for *C. difficile* infection. Upon their arrival, fecal pellets were collected from all hamsters, homogenized in 1 mL saline, and examined for *C. difficile* by plating on CCFA-TA (cycloserine cefoxitin fructose agar-0.1% taurocholate) to ensure that the animals did not harbor endogenous *C. difficile*. After this initial screen, they were housed individually in sterile cages with ad libitum access to food and water for the duration of the study. Hamsters were first gavaged with 30 mg/kg clindamycin(Hasan *et al*., 2024a). Infection was initiated 3 days after clindamycin administration by oral gavage with 2000 *C. difficile* spores. Strains UK1 (parent with *codY*(WT), UK1*codY*::*ermB* (*codY* mutant), and strains expressing *codY* variants, UK1::*codY*(V58A) and UK1:: *codY*(Y146N) were used in this experiment. Ten animals per group, including an uninfected control group (with six animals), were used in the studies. Colonization was monitored by plating fresh fecal pellets in a *C. difficile* selective medium. Percentage survival was recorded as described previously (Girinathan *et al*., 2018; Hasan *et al*., 2024). Briefly, animals were monitored for signs of disease (lethargy, poor fur coat, sunken eyes, hunched posture, and wet tail) every 4 hours (six times per day) throughout the study period. Hamsters were scored from 1 to 5 for the signs mentioned above (1, normal, and 5, severe). Hamsters showing signs of severe disease (a cumulative score of 12 or above) were euthanized by CO_2_ asphyxiation. Surviving hamsters were euthanized 15 days after *C. difficile* infection. The survival data of the challenged animals were graphed as Kaplan-Meier survival analyses and compared with statistical significance using the log-rank test using GraphPad Prism.

## Results

### Naturally occurring CodY variants

CodY is a nutrient-sensing master regulator in several pathogens as well as model organisms such as *B. subtilis* (Ratnayake-Lecamwasam *et al*., 2001; Shivers and Sonenshein, 2004; Sonenshein, 2007). Analysis of amino acid sequences among available sequenced *C. difficile* strains shows the existence of several versions of CodY. These include alleles with the following mutations: T18K, A46S, L55M, V58A, E89Q, Y146N, S145C, and G152S. Among these, we chose CodY(Y146N) and CodY(V58A) alleles for further analysis since the altered residues (Y146 and V58) are close to the known ligand-binding sites in CodY (Fig. 1A and 1B). The Y146N mutation is present in the R20291 derivative strain. Laboratories worldwide commonly use the *C. difficile* R20291 as a model strain to study the highly virulent NAP027 ribotype. Recently, through whole genome sequencing, it was found that three different R20291 stocks with various point mutations exist. The strain we use in our lab was originally obtained from Dr. Trevor Lawley’s lab at the Sanger Institute. This strain (R20291-CRG 3661) carries the Y146N mutation in the *codY* gene (Monteford *et al*., 2021). The *codY(*V58A) allele is the second most common *codY* allele in *C. difficile* strains and is present in different clinically relevant ribotypes. For example, the strain M120 belonging to the ribotype 078 carries the CodY(V58A) allele. The ribotype 078 is widespread among swine and cattle and is an emerging strain in humans (Smits, 2013; Aboutaleb *et al*., 2014; Kachrimanidou *et al*., 2019). Characterizing the impact of CodY V58A and Y146N variants can shed light on how point mutations in an important master regulator could change the pathophysiology of a pathogen.

**Figure 1.**
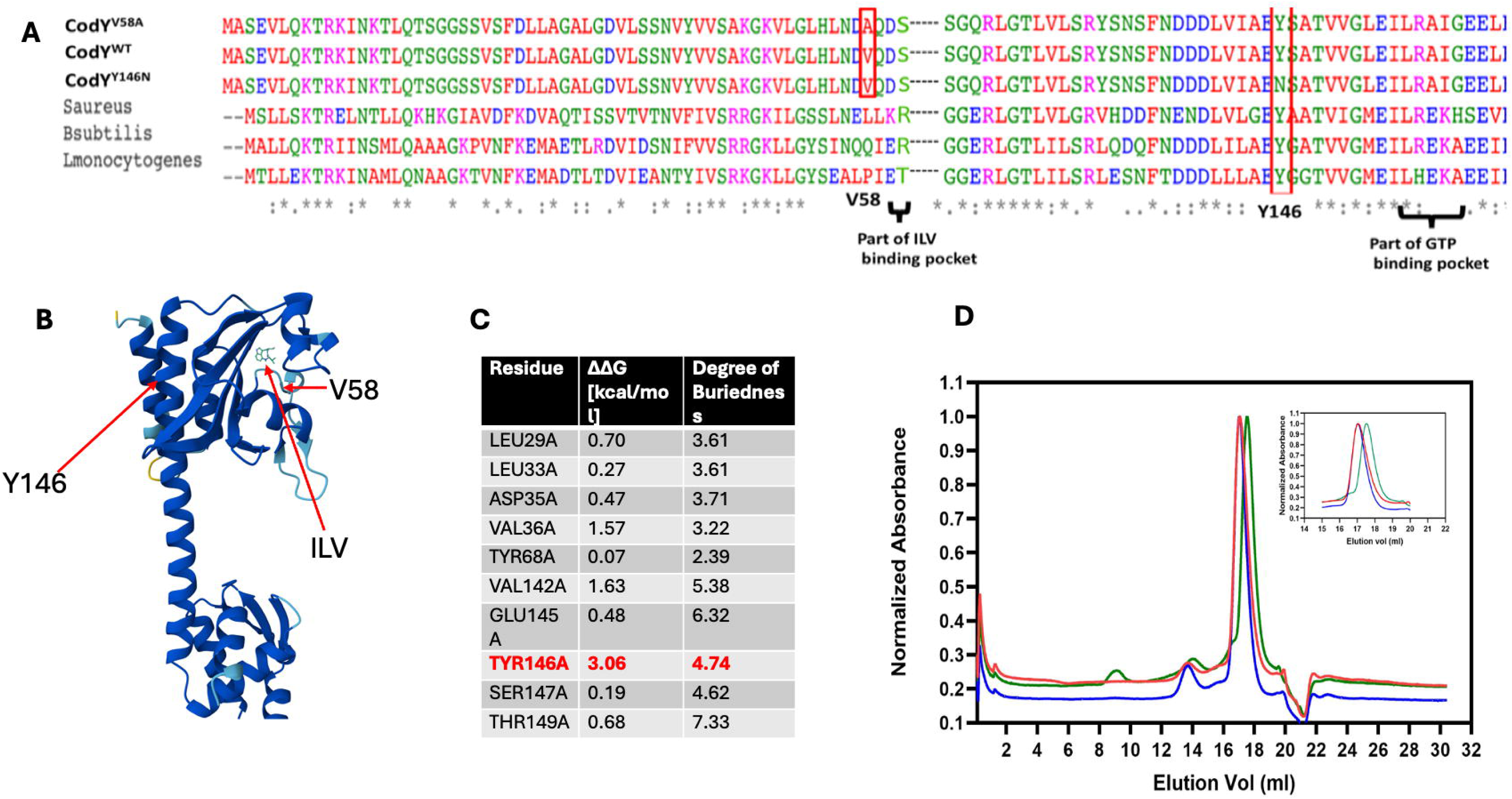
**A**. Clustal-W alignment of *C. difficile* CodY variants with CodY in different Gram+ bacteria. V58 and Y146 residues are highlighted in the red box. Residues involved in ILV and GTP binding are also indicated. **B.** AlphaFold structure of full-length *C. difficile* CodY with isoleucine as ligand. Positions of V58 and Y146 residues are indicated. **C.** Alanine scanning results of CodY residues near V58 and Y146. **D.** Analytical gel filtration runs of CodY(WT) (red), V58A (blue), and Y146N (green) run on Superdex™ 200 Increase 10/300 GL column (Cytiva) with normalized absorbances, with inset magnifications to highlight elution volume difference.

### Mutation of Y146 disrupts CodY dimerization as predicted by *in silico* analysis

CodY is recognized for its ability to link nutrient availability, metabolism, and virulence gene expression in Gram-positive pathogens such as *C. difficile* and *Staphylococcus aureus* by sensing branched-chain amino acids-ILV and GTP (Pohl *et al*., 2009; Waters *et al*., 2016).

Crystal structures of full-length CodY from *Bacillus subtilis*, *S. aureus*, *Bacillus cereus*, and *Enterococcus faecalis* have been resolved (Han *et al*., 2016; Levdikov *et al*., 2017). However, only the truncated form of *C. difficile* CodY, consisting of the first 155 residues, has been resolved by crystal structure (Fig. 1B) (Daou *et al*., 2019a). Since our point mutants of interest are encompassed within these residues, we used this structure (PDB 5N0L) for our analysis. The residue V58, exclusive to *C. difficile* CodY, is located near the ILV-binding pocket (Fig. 1A, 1B), and any modification to this residue could impact ILV binding. The residue Y146, situated at the dimer interface, is highly conserved among CodY proteins in other Gram-positive bacteria (Fig. 1A). To identify residues that are significant for CodY dimerization, we used the DrugScorePPI web tool (Krüger and Gohlke, 2010) to perform in silico alanine scanning mutagenesis (supplementary method) and measure the change in binding free energy (ΔΔG) of CodY dimers. The program swaps each residue of CodY to Alanine to test whether the mutation will create a more or less stable dimer, which is expressed as a measurement of binding free energy (ΔΔG). Consequently, a higher ΔΔG value indicates the mutation will destabilize the dimer formation, whereas negative values indicate the mutation will further stabilize the dimer. Unchanged values (ΔΔG =0) indicate that the residue has no effect on dimerization or is not located at the dimerization interface. The analysis identifies several residues in the dimer interface with significant effects on dimer stability, such as V36, V142, and T149. However, among the identified residues, mutation of Y146 leads to the highest predicted perturbation in the dimerization potential of CodY (ΔΔG of 3.06) (Fig. 1C).

CodY dimerization is essential for its proper transcriptional regulation (Joseph *et al*., 2005; Han *et al*., 2016). To validate the *in silico* predictions regarding dimerization-disrupting mutations, we analyzed the oligomeric state of purified WT and mutant CodY proteins by size-exclusion chromatography (SEC). For each analytical gel filtration run, 500 μL of protein sample was injected into a Superdex™ 200 Increase 10/300 GL column (Cytiva), and the elution profiles were monitored at 280 nm. The wild-type CodY protein eluted at 17.06 mL, the V58A mutant at 17.03 mL, and the Y146N mutant at 17.55 mL (Fig. 1D). In SEC, larger biomolecules or complexes elute earlier than smaller ones, and the apparent molecular weight is estimated by comparing the elution volume with that of protein standards of known size. Based on the elution volumes, the apparent molecular weights of the WT and V58A CodY proteins were estimated to be 51.8 and 53.2 kDa, respectively, whereas the Y146N mutant had an apparent molecular weight of 34.1 kDa. Considering that the predicted molecular weight of recombinant CodY (including the His-tag and TEV recognition sequence) is approximately 30 kDa, these results indicate that WT and V58A CodY behave predominantly as dimers (expected dimeric size ∼60 kDa), while the Y146N mutant exists mainly as a monomer. The apparent underestimation of molecular weight for the dimeric forms (51.8–53.2 kDa vs. 60 kDa) likely arises from differences in molecular shape between CodY and the globular proteins used as SEC standards. Unlike the spherical standards, full-length CodY adopts an elongated, dumbbell-like conformation (Fig. 2B), which can alter its hydrodynamic radius and lead to a smaller apparent molecular weight in SEC analysis.

**Figure 2.**
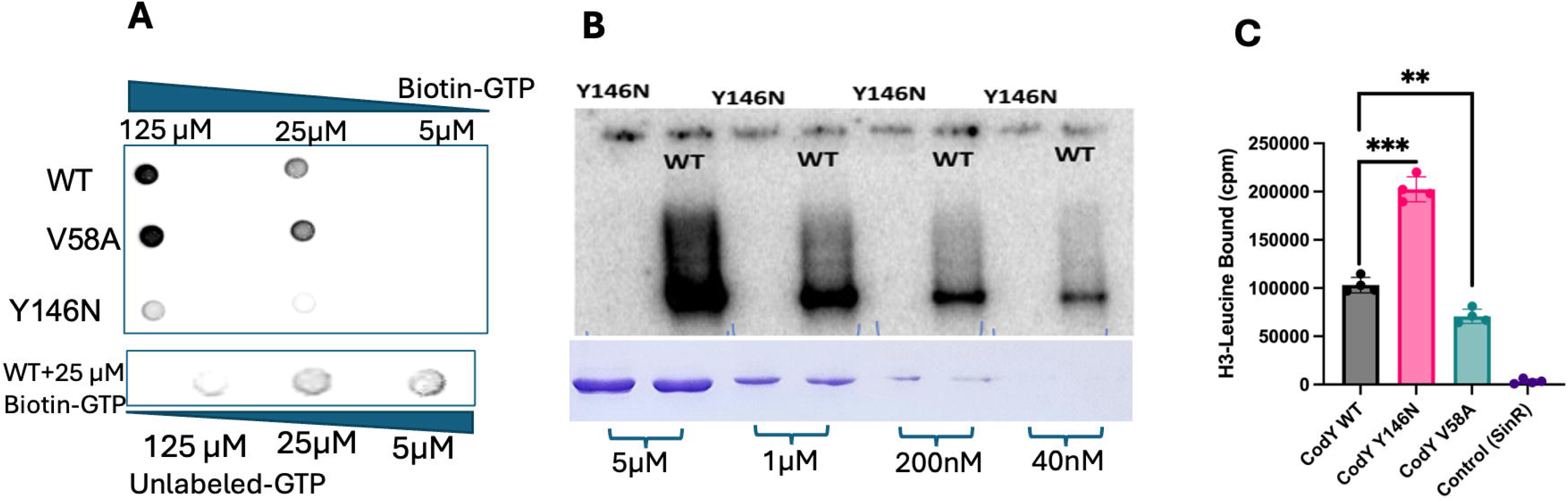
A. CodY binding assay with purified CodY. Purified CodY proteins (50 µM) were incubated with biotin-labeled GTP and UV cross-linked. Reaction mixtures were spotted onto a PVDF membrane and were developed after incubation with Streptavidin-HRP conjugate (Top panel). Biotin-GTP binding specificity was tested by the addition of unlabeled GTP (Lower panel). **B.** Purified CodY proteins were incubated with [γ-³²P]GTP, resolved by SDS-PAGE, and visualized using a Typhoon phosphorimager. A parallel gel loaded with the same amount of protein but without [γ-³²P]GTP was stained with Coomassie to compare protein quantities between CodY(WT) and CodY(Y146N) (lower panel). **C.** Leucine binding activity tested with purified CodY proteins incubated with [H3]-Leucine, followed by UV cross-linking. Proteins in the reaction were TCA precipitated, and the radioactivity associated was measured by scintillation counter. Data represent mean ± SEM from three independent experiments. Statistical significance was determined by one-way ANOVA followed by Dunnett’s post hoc test comparing each mutant to the wild type (*P* < 0.05). Purified SinR protein used as a negative control.

### CodY-Y146N failed to bind to GTP

In order to test GTP binding, 30 µM of His-tagged CodY(Y146N), CodY(V58A), and CodYWT were purified and incubated with various concentrations of biotin-labeled GTP (NEB biolabs). After UV-cross-linking the binding reactions, the proteins were spotted onto the PVDF membrane and probed with streptavidin-HRP. The results showed that only CodY(WT) and CodYV58A exhibited binding to GTP, while CodY(Y146N) showed very little affinity to GTP (Fig. 2A). To check for the specificity of this binding, we incubated different concentrations of unlabeled GTP with the CodY-WT (Fig. 2A). In this competition binding reaction, with increasing concentration of unlabeled GTP, less biotin-GTP was detected binding to the CodY protein. To further verify the GTP-binding deficiency of CodY(Y146N), a more sensitive binding assay with radiolabeled [γ-³²P] GTP was performed. Binding reactions containing various amounts of CodY(WT) or CodY(Y146N) were incubated with 1 µCi of [γ-³²P] GTP (∼0.125 µM) in a 10 µL reaction prior to cross-linking. The samples were then separated by SDS–PAGE and visualized using a Typhoon phosphor imager. Because the concentration of [γ-³²P] GTP was approximately 1,000-fold lower than that used in the biotin–GTP assay, only CodY(WT) showed detectable binding, even at low protein concentrations (Fig. 2B). These results demonstrate that the Y146N substitution severely impairs GTP binding, likely due to altered dimerization and/or structural disruption of the ligand-binding domain.

To evaluate leucine binding, the purified CodY(WT) and variant proteins were incubated with radiolabeled [³H]-leucine, and the bound ligand was quantified by scintillation counting. All three CodY proteins retained the ability to bind leucine but exhibited distinct binding affinities. CodY(Y146N) showed nearly twofold higher binding than CodY(WT), whereas CodY(V58A) exhibited moderately reduced binding compared to the wild type (Fig. 2C). These findings indicate that, while the Y146N mutation disrupts GTP binding, it does not abolish leucine recognition and may even enhance it, possibly due to increased accessibility to leucine.

### Differential binding of CodY variants to the target gene promoter

The binding of CodY to its ligands is known to influence its affinity for target DNA. Since we observed differences in ligand binding among the CodY variants, we next examined their ability to bind one of CodY’s target DNAs, the *tcdR* upstream region. The *tcdR* gene encodes an alternative sigma factor that positively regulates toxin gene expression, whereas CodY represses toxin production by binding to P*tcdR* and inhibiting *tcdR* transcription.

Radiolabeled DNA fragments containing P*tcdR* were used in EMSAs with CodY variants, and binding was assessed in the presence or absence of ligands (1 mM GTP, 10 mM ILV, or both) (Fig. 3). In the absence of ligands, a DNA shift was observed at a minimal CodY concentration of 63 nM for all three proteins. CodY(WT) exhibited increased affinity for P*tcdR* in the presence of ILV or GTP, as DNA binding was detectable at lower protein concentrations (8 nM and 31 nM, respectively). In contrast, the CodY(Y146N) and CodY(V58A) proteins did not show enhanced binding to P*tcdR* in the presence of either ligand (Fig. 3). These results suggest that ILV binding does not enhance the DNA-binding activity of CodY(Y146N), despite its higher affinity for ILV and possibly due to its reduced ability to form functional dimers. This result suggests that differential binding of CodY variants to their target DNAs could lead to altered transcriptional regulation of these genes, ultimately affecting the virulence potential of the strains producing them. To test this possibility, we next examined the impact of CodY variants on *C. difficile* virulence.

**Figure 3.**
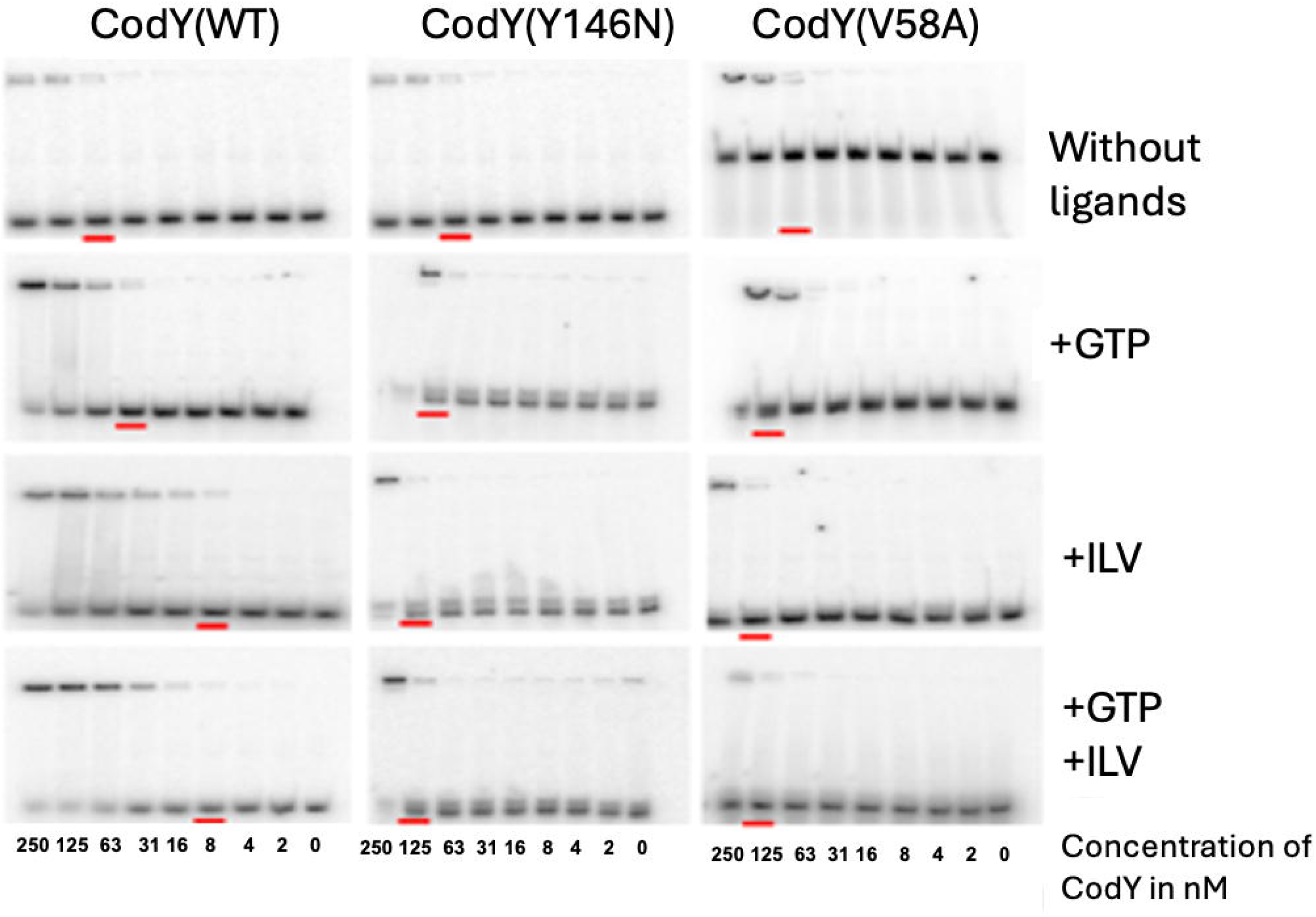
EMSA performed with purified CodY (WT) or CodY(Y146N), or CodY(V58A) with the *tcdR* promoter. ILV or GTP or both were added as indicated. The lowest concentration of protein needed for a visible shift is underlined in red.

### Effect of CodY variants on toxin production in *C. difficile*

CodY represses toxin production in *C. difficile* by modulating *tcdR* transcription. Since we observed differences in the binding of CodY variants to P*tcdR*, we next investigated whether these CodY mutations influence toxin production. To directly assess the effect of CodY variants in an identical genetic background, we introduced *codY*(Y146N) and *codY*(V58A) alleles into the *codY* mutant (UK1*codY::ermB*) of the UK1 strain (Daou *et al*., 2019a) using the pMSR0-based allele exchange method (Peltier *et al*., 2020). This strategy ensured that the only variable among the strains was the *codY* allele itself.

The *codY* variants were integrated into the native chromosomal locus, replacing the intron-disrupted *codY* in the mutant strain. Growth analyses revealed no significant differences among the strains expressing CodY(WT), CodY(Y146N), or CodY(V58A) under the tested conditions (Fig. S1). Western blot analysis confirmed production of CodY variants prior to assessing toxin production (Fig. 4A). Cytosolic toxin levels were measured in bacteria grown under various conditions (without ligands, with 100 μM guanosine, or with 10 mM ILV). Guanosine was added to the medium since it serves as the precursor for GTP biosynthesis (King *et al*., 2018a). As shown in Figure 4A, both CodY(V58A) and CodY(Y146N) were markedly less efficient in repressing toxin production compared to CodY(WT) under all tested conditions. The addition of ILV or GTP did not enhance the repressive activity of either mutant protein. Consistent with previous findings, CodY(WT) displayed strong repression of toxin production in the presence of ILV (Dineen *et al*., 2007; Dineen *et al*., 2010), whereas the CodY variants failed to respond to the ligands. These results indicate that the altered ligand- and DNA-binding properties of CodY(V58A) and CodY(Y146N) impair their ability to regulate toxin synthesis in *C. difficile*.

**Figure 4.**
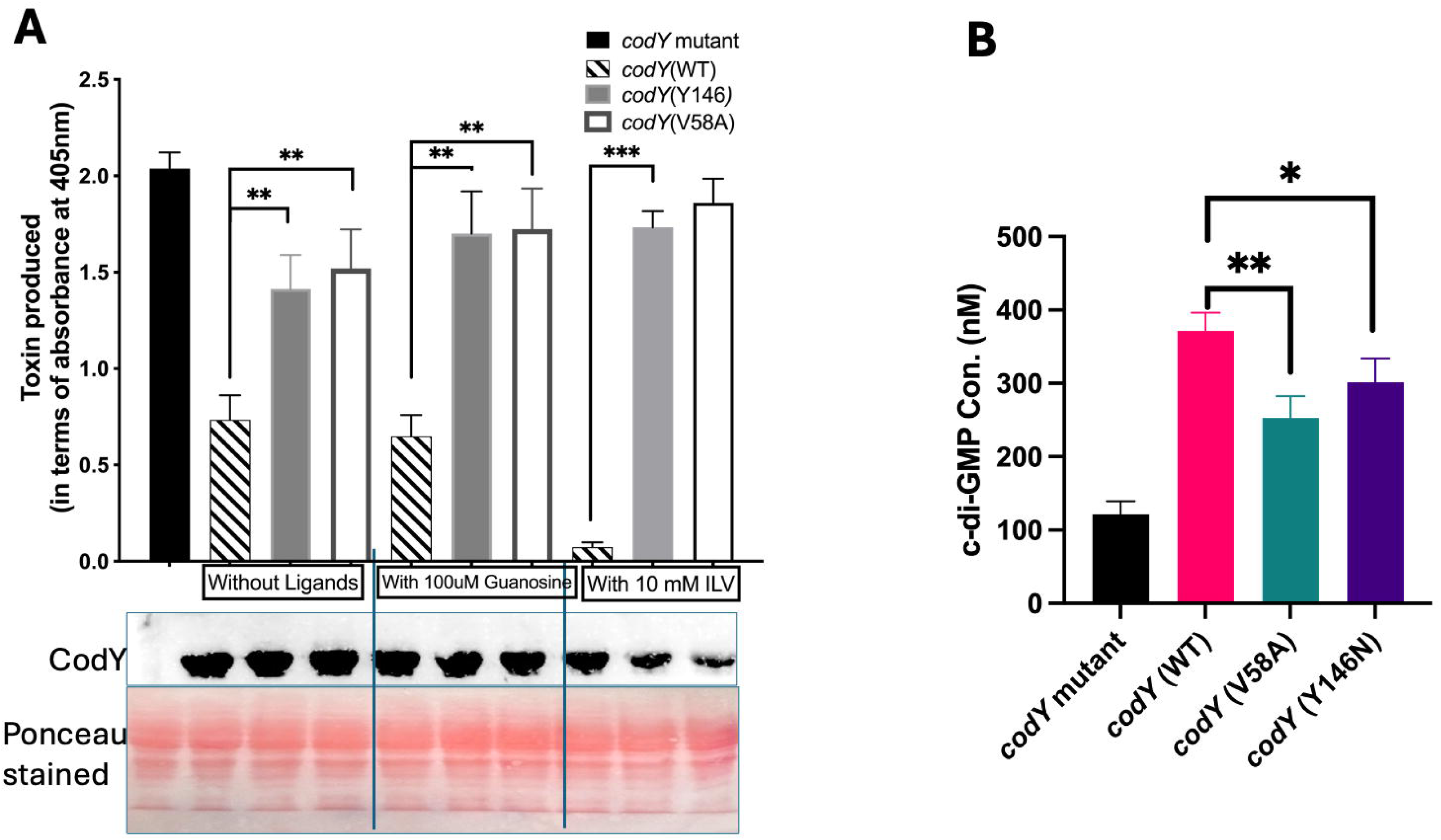
A. Effect of *codY* alleles on toxin production *in C. difficile*. Toxin production (measured using ELISA) in *codY mutant* (UK1*codY::ermB*) and *C. difficile* strains producing CodY (WT) or CodY (Y146N), or CodY (V58A). CodY production by these strains were confirmed by western blot with anti-CodY antibodies. The membrane was stained with Ponceau stain to show equal loading of proteins. **B.** Measurement of intracellular levels of c-di-GMP in *C. difficile* cytosol using the c-di-GMP Assay kit (Lucerna). Concentrations were calculated using a standard curve constructed using known amounts of c-di-GMP, and the data presented indicate the c-di-GMP concentration in the volume (20µl) of cytosol tested. Data represent mean ± SEM from three independent experiments. Statistical significance was determined by one-way ANOVA followed by Tukey’s multiple-comparison test (* p < 0.05; ** p < 0.01; *** p < 0.001).

### Indirect regulation of toxin gene expression by CodY through c-di-GMP signaling

In addition to its direct regulation of toxin genes via *tcdR*, CodY also indirectly influences toxin expression by modulating intracellular levels of cyclic di-GMP (c-di-GMP), a key signaling molecule that controls motility and toxin gene expression in *C. difficile*. c-di-GMP binds to a riboswitch located upstream of the flagellar operon and suppresses its transcription (El Meouche *et al*., 2013). The flagellar operon includes the gene encoding the sigma factor SigD, which positively regulates *tcdR* transcription and, consequently, toxin gene expression (McKee *et al*., 2013). Therefore, elevated intracellular c-di-GMP levels lead to reduced toxin production, while lower c-di-GMP levels promote toxin synthesis. Intracellular c-di-GMP levels are determined by the balance between diguanylate cyclases, which synthesize c-di-GMP, and phosphodiesterases (PDEs), which degrade it. CodY represses the PDE-encoding genes *pdcB* and *pdcA* (Purcell *et al*., 2017; Dhungel and Govind, 2021). When CodY repression is active, PDE expression is reduced, leading to c-di-GMP accumulation. Conversely, weakened CodY repression increases PDE production, resulting in enhanced c-di-GMP degradation, elevated *sigD* expression, and subsequently higher *tcdR*, *tcdA*, and *tcdB* transcription, resulting in increased toxin levels.

To determine whether CodY variants alter intracellular c-di-GMP concentrations, we quantified c-di-GMP levels in the cytosolic fractions of the corresponding strains. As expected, strains expressing CodY(Y146N) and CodY(V58A) contained significantly lower levels of c-di-GMP compared to those producing CodY(WT) (Fig. 4B). Collectively, these results demonstrate that mutations in critical functional domains of this master regulator can reshape the expression of its target genes and, in turn, could modulate *C. difficile* pathogenesis and virulence.

### *C. difficile* expressing *codY* variants are highly virulent

The *C. difficile* hamster infection model is a well-established system for evaluating the pathogenic potential of *C. difficile* strains, as it closely mimics the clinical progression of human disease (Buckley *et al*., 2011; Weiss *et al*., 2014). Using this model, we assessed the virulence potential of *C. difficile* strains expressing CodY variants. Within 30 hours post-challenge, animals infected with strains producing CodY(Y146N) or CodY(V58A) showed efficient colonization, exhibiting significantly higher CFU counts in fecal samples than those infected with the CodY(WT)-producing strain (Fig. 5A). The codY mutant (UK1 *codY*::*ermB*) was excluded from colonization analysis because animals infected with this strain developed severe diarrhea and succumbed rapidly to infection (Fig. 5B), consistent with the hypervirulent phenotype previously associated with *codY* disruption.

**Figure 5.**
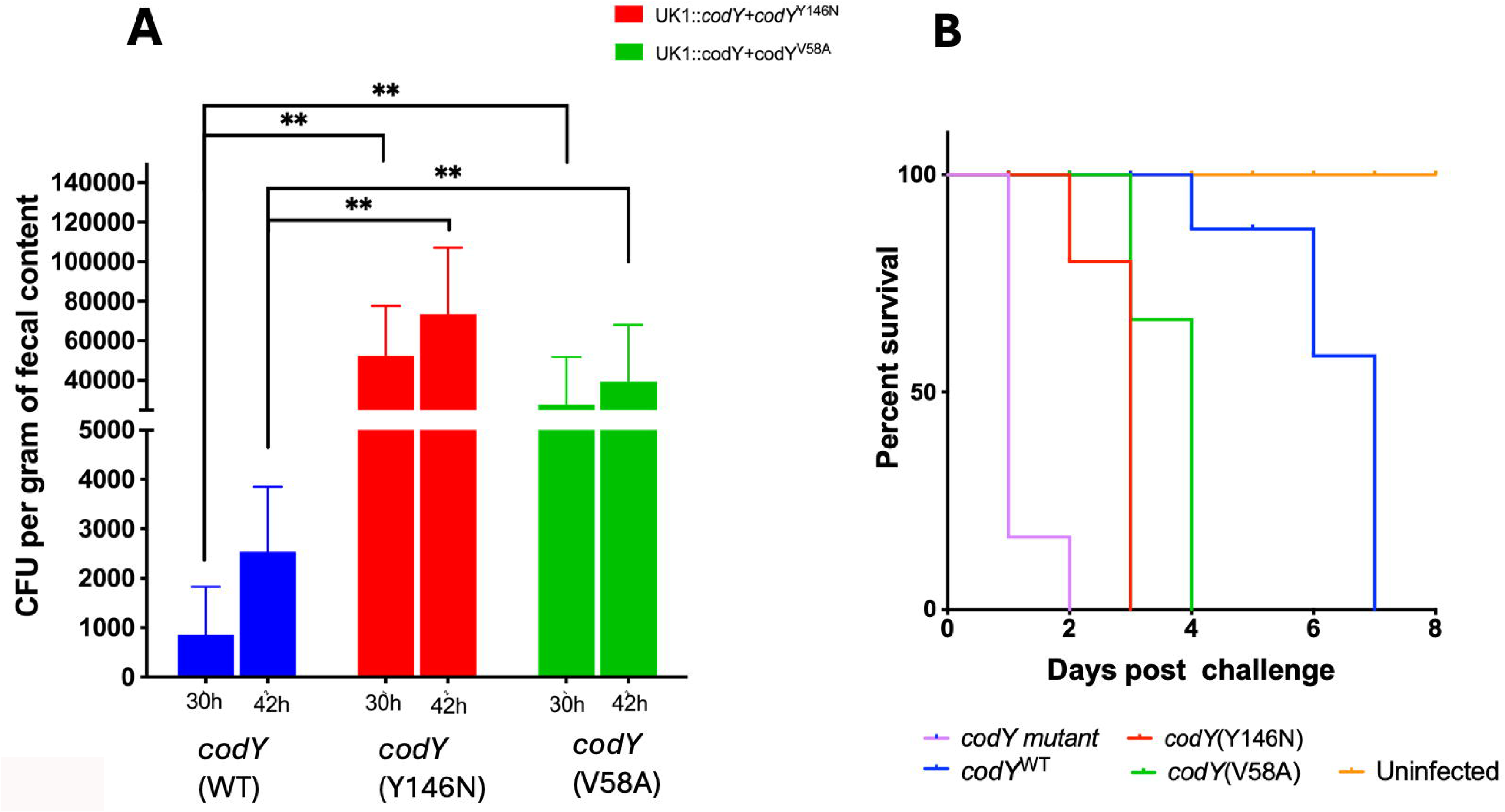
A. Colonization capacity of different CodY variants compared at 30h and 42h after challenge by enumerating CFU of *C. difficile* recovered from fecal samples. Data was analyzed by one-way analysis of variance (ANOVA), where *** indicates p<0.001. **B.** Kaplan-Meier survival plot of Hamster groups infected with *codY mutant or C. difficile* strains expressing either *codY* WT or *codY* (Y146N), or *codY* (V58A). Survival curves were analyzed using the Kaplan– Meier method, and significance was determined by the log-rank (Mantel–Cox) test (*P* < 0.05).

Survival analysis further confirmed enhanced virulence associated with the CodY variants (Fig. 5B). Animals challenged with strains expressing CodY(Y146N) or CodY(V58A) exhibited significantly reduced survival compared to those infected with the CodY(WT)-producing strain. Collectively, these findings are consistent with our in vitro observations and demonstrate that naturally occurring mutations in CodY can increase *C. difficile* virulence, likely through impaired regulation of virulence-associated pathways, including toxin production.

## Discussion

Ligand-binding transcriptional master regulators act as central hubs that integrate nutrient sensing and environmental cues to control virulence gene expression. Mutations that alter their ligand-binding or allosteric properties can profoundly affect pathogenicity. In *Listeria monocytogenes*, for example, a naturally occurring G145S mutation in PrfA master regulator stabilizes its active conformation, producing a constitutively active, hypervirulent phenotype. Similarly, L47P and E178K substitutions in *Mycobacterium bovis* CRP abolish cAMP-dependent activation while preserving ligand binding (Gárate *et al*., 2021). These examples underscore how subtle amino acid changes can remodel global regulatory networks governing virulence.

Our study identifies two variants of the nutrient-sensing regulator CodY that rewire virulence control in *C. difficile*. CodY functions as a metabolic gatekeeper, responding to GTP and branched-chain amino acids (BCAAs) to repress virulence-associated genes (Dineen *et al*., 2010; King *et al*., 2018; Daou *et al*., 2019). Altered ligand sensing by these variants disrupts this balance. CodY(Y146N) showed reduced affinity for GTP but enhanced binding to leucine, while CodY(V58A) exhibited the opposite trend. The Y146N mutation impaired CodY dimerization, a prerequisite for full activity, consistent with alanine scanning and SEC analyses. Y146 is highly conserved across CodY orthologs (Fig. 1A), suggesting a structurally critical role that likely extends to other Gram-positive bacteria. We performed *in silico* alanine mutagenesis using B. subtilis CodY and confirmed that mutation of Y145 (residue equivalent to Y146) would likely lead to similar disruption in dimerization (S3 Table).

Beyond altered ligand binding, both CodY variants exhibited defective ligand-dependent DNA binding, indicating disrupted allosteric communication between the effector- and DNA-binding domains. Both the CodY variants bound weakly to the *tcdR* promoter, which could explain the elevated toxin levels in the strains producing CodY(Y146N) or CodY(V58A). Reduced intracellular c-di-GMP levels recorded in these strains, likely through the derepression of the phosphodiesterases coding genes *pdcA* and *pdcB,* could also result in elevated *sigD* and *tcdR* expression, which in turn can increase toxin production. Increased colonization and lethality of CodY(Y146N) and CodY(V58A) producing strains in the hamster model further confirmed the effect of CodY modifications to enhanced virulence. Analogous single-point mutations that modify bacterial virulence have been reported in other systems, for instance, the Y223C substitution in *Staphylococcus aureus* AgrC, which enhances dimerization but weakens interaction with AgrA, leading to hyperactivation of virulence genes (Mairpady Shambat *et al*., 2016). These findings demonstrate that single amino acid substitutions in global regulators can relax normal constraints on virulence gene expression and reshape host-pathogen interactions.

It is important to note that the CodY(Y146N) variant is present in a commonly used R20291 laboratory stock (Monteford *et al*., 2021), and results from studies using this strain should therefore be interpreted with caution. The CodY(V58A), located near the ILV-binding pocket, is prevalent in ribotype 078 strains linked to livestock and severe disease, and could contribute to their elevated virulence. From an evolutionary standpoint, CodY variants exemplify how regulatory evolution drives pathogen diversification. Rather than acquiring new virulence genes, *C. difficile* can increase pathogenic potential through subtle modifications in global regulatory hubs. Monitoring such naturally occurring polymorphisms in CodY and related regulators may offer early insights into the emergence of hypervirulent lineages.

